# Ensemble modeling of auditory streaming reveals potential sources of bistability across the perceptual hierarchy

**DOI:** 10.1101/598631

**Authors:** David F. Little, Joel S. Snyder, Mounya Elhilali

## Abstract

Perceptual bistability—the spontaneous fluctuation of perception between two interpretations of a stimulus—occurs when observing a large variety of ambiguous stimulus configurations. This phenomenon has the potential to serve as a tool for, among other things, understanding how function varies across individuals due to the large individual differences that manifest during perceptual bistability. Yet it remains difficult to interpret the functional processes at work, without knowing where bistability arises during perception. In this study we explore the hypothesis that bistability originates from multiple sources distributed across the perceptual hierarchy. We develop a hierarchical model of auditory processing comprised of three distinct levels: a Peripheral, tonotopic analysis, a Central analysis computing features found more centrally in the auditory system, and an Object analysis, where sounds are segmented into different streams. We model bistable perception within this system by injecting adaptation, inhibition and noise into one or all of the three levels of the hierarchy. We evaluate a large ensemble of variations of this hierarchical model, where each model has a different configuration of adaptation, inhibition and noise. This approach avoids the assumption that a single configuration must be invoked to explain the data. Each model is evaluated based on its ability to replicate two hallmarks of bistability during auditory streaming: the selectivity of bistability to specific stimulus configurations, and the characteristic log-normal pattern of perceptual switches. Consistent with a distributed origin, a broad range of model parameters across this hierarchy lead to a plausible form of perceptual bistability. The ensemble also appears to predict that greater individual variation in adaptation and inhibition occurs in later stages of perceptual processing.

**Author summary:** Our ability to experience the everyday world through our senses requires that we resolve numerous ambiguities present in the physical evidence available. This is accomplished, in part, through a series of hierarchical computations, in which stimulus interpretations grow increasingly abstract. Our ability to resolve ambiguity does not always succeed, such as during optical illusions. In this study, we examine a form of perceptual ambiguity called bistability—cases in which a single individual’s perception spontaneously switches back and forth between two interpretations of a single stimulus. A challenge in understanding bistability is that we don’t know where along the perceptual hierarchy it is generated. Here we test the idea that there are multiple origins by building a simulation of the auditory system. Consistent with a multi-source account of bistability, this simulation accurately predicts perception of a simple auditory stimulus when bistability originates from a number of different sources within the model. The data also indicate that individual differences during ambiguous perception may primarily originate from higher levels of the perceptual hierarchy. This result provides a clue for future work aiming to determine how auditory function differs across individual brains.

## Introduction

Perceptual bistability—the spontaneous fluctuation of perception between two interpretations of a stimulus—can occur while observing ambiguous stimulus configurations [1–7]. A classic example is the Necker cube [1]. This image, comprised of an abstract line-drawing of a transparent 3D cube, can be interpreted as a cube seen from above or from below. If an individual stares at the cube, over time, perception will fluctuate between the above- and the below-view interpretations. In the auditory system, a prominent example of bistability comes from the classic auditory streaming paradigm [8, 9], in which a repeating A-B-A pattern of pure tones appears to be one (ABA) or two (A-A, and -B-) objects [4]. Here, we test the idea that perceptual bistability is generated from multiple sources distributed throughout the brain [3, 10–20], using a large ensemble of computational models of this auditory streaming paradigm.

The study of perceptual bistability, and the more general phenomena of multistability, has the potential to shed light on a number of fundamental principles of perception, as evidenced by four properties. First, its manifestation is characterized by substantial individual variation [21–29] providing a potential means for understanding how perceptual function differs across individuals. Second, it is a quite general phenomenon, as there are many forms of ambiguous stimuli that can generate perceptual bistability or multistability [1–4, 6]. These include not just abstract, laboratory stimuli, but also complex stimuli such as speech [6] and faces [3]. Third, the proposed mechanisms of multistability are also candidate mechanisms for decision making [18, 30–32] and perceptual inference [5, 7, 18]. Fourth, bistability provides a case in which perception varies while the observed stimulus remains constant, a valuable control when testing theories of attention, awareness and consciousness [33–36].

A number of properties of bistability can be accounted for using three simple ingredients: adaptation, inhibition and noise [18, 19, 37–42, 42–45]. In isolation, inhibition can be used to define a set of attractors of neural activity; this in turn can lead to a winner-take-all behavior. Activity can shift from one of these attractors to another due either to noise or due to adaptation of the winning attractor. Appropriate forms of these dynamics can in turn be used to implement a number of fundamental computational elements necessary for decision making [18, 30–32] and perceptual inference [5, 7, 18]. While these ingredients appear to reflect quite fundamental neural-computational principles, the actual impact on perception more broadly cannot be determined without knowing where bistability arises during perception.

An emerging idea is that, rather than originating from a single source, perceptual bistability is driven by many different sources of adaptation, inhibition and noise across the brain, in a distributed fashion [3, 10–20]. This idea has largely been invoked to account for an apparent contradiction in the literature: there is evidence which appears to favor an early locus [46–50], whereas other evidence suggests a much later locus [51–58] for perceptual bistability. A natural resolution to these differing outcomes is that there are multiple, distributed sources of bistability [3, 11–17]. Another source of evidence for this distributed account comes from a careful examination of the distribution of reported percept lengths—the pattern of switches from one percept to another—which appear to be best explained by multiple bistable sources [19, 20].

If perceptual bistability really emerges from multiple sources across the perceptual hierarchy, then it should be possible to induce a plausible form of perceptual bistability within a model of this hierarchy. Yet, past efforts to model perceptual bistability do not systematically vary the locus and the magnitude of adaptation, inhibition and noise within a perceptual hierarchy. This makes it difficult to asses the relative merits of different bistable loci within that perceptual hierarchy, as there are many confounding differences across computational models. Most mathematical models have focused on a single, abstract level of processing upon which adaptation, inhibition and noise generate an oscillatory response [18, 37–43, 45, 59–61]. The most relevant efforts to date, which include multiple hierarchical levels of processing [11, 13, 14, 17, 36], do not systematically vary the configuration of multiple loci of adaptation, inhibition and noise. Those efforts that do examine systematic variation of the parameter space [15, 18, 19, 36, 43, 45] consider only a single locus for bistability. An important reason for these limitations is that these past models have generally employed a relatively simplified and abstract description of perceptual input.

To begin to address these limitations, in the present report we evaluate a series of detailed hierarchical models of human behavior during a simple bistable auditory streaming task [4, 8, 9]. The model includes three hierarchical stages: *Peripheral*, *Central* and *Object*. The Peripheral stage performs a time-frequency analysis; this is followed by the Central analysis which computes spectral features found more centrally in the auditory system. In the third, Object analysis, these features are bound into a probabilistic interpretation of the acoustic streams (or objects) present in the scene. Each model can be understood as some variant of an abstraction of the ventral auditory system [62, 63], in the sense that it generates an interpretation of what sources are present in an auditory scene, and does so in manner consistent with principles of scene analysis gleaned from past empirical data [8, 9, 64–66].

Across the three stages of this hierarchy, we systematically vary adaptation, inhibition and noise within a model ensemble. Ensemble modeling comprises the evaluation of many different mathematical models, each varying in their parameters and/or configuration. With this large variety of parameter configurations, some of the models perform poorly, failing to reflect human behavior, but a subset provide a good fit to the empirical data. This approach avoids the assumption that a single mechanism—represented by a single parameter configuration—must be invoked to explain all of the experimental data [67, 68]. Rather, there may be many models within the ensemble capable of accounting for the available data, reflecting variation that may occur both within and across individuals.

When we systematically vary the level of adaptation, inhibition and noise across the auditory hierarchy, we find that a plausible form of bistability can be generated using these three ingredients at any one of the three different stages of analysis or across all stages simultaneously. The results are therefore consistent with a multi-source hypothesis for perceptual bistability. Furthermore, the range of levels of adaptation and inhibition that generate a plausible form of bistability is much larger at the highest stage of this hierarchy (Object), while a much more precise tuning of adaptation and inhibition is required at the earliest stage of analysis (Peripheral)—and the Central-analysis parameter tuning is somewhere in-between these extremes. These differences in parameter sensitivity across the model hierarchy raise an interesting possibility, given the evidence for considerable variability across listeners during bistable perception [21–29]. It may be that these individual differences across human listeners originate primarily from later stages of the perceptual hierarchy, while earlier stages have a more precise tuning of adaptation and inhibition.

## Results

### Model Description

Fig 1A shows our computational framework: it includes three levels of processing: Peripheral (Fig 1A; left panel), Central (Fig 1A; middle panel) and Object (Fig 1A; right panel). Within each of these levels, we inject a given amount of adaptation, inhibition and noise, shown in Fig 1B. Full details of the model design can be found in Materials and methods.

**Fig 1.**
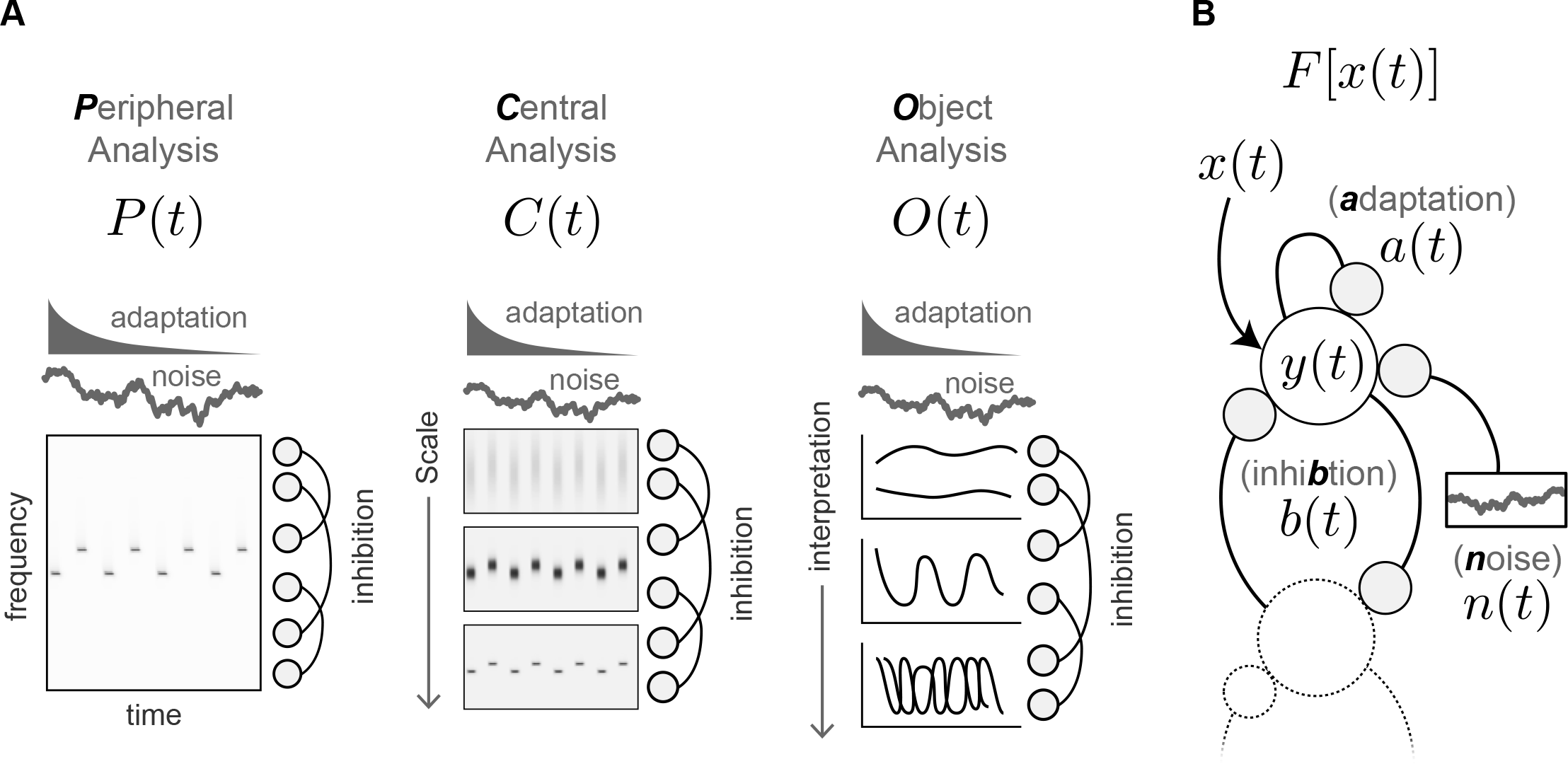
Model Design. Each model in the ensemble includes three hierarchical analysis stages: Peripheral, *P* (*t*), Central, *C*(*t*), and Object, *O*(*t*), into which we inject adaptation, inhibition and a small amount of noise, *F* [*x*(*t*)] **A**. The three analysis stages. The Peripheral analysis, *P* (*t*), computes a log-frequency spectrum using cochlear-like filter shapes. The Central analysis, *C*(*t*), computes multiple spectral scales of *P* (*t*), to capture dependencies across different frequencies. The Object analysis, *O*(*t*), computes multiple interpretations of the auditory scene into separate masks of *C*(*t*), selecting the interpretation most consistent with the data at each time frame. **B**. The application of adaptation, inhibition and noise. Adaptation, *a*(*t*), reduces the input, *x*(*t*), by a low pass version of the output history; inhibition, *b*(*t*), reduces the input by a low pass version of the output history of distant neighbors; noise, *n*(*t*), adds a small amount of variation to the output weights. The result is a set of output weights, *y*(*t*), applied within each model along the frequencies (for Peripheral), scales (for Central) or scene interpretations (for Object).

During the Peripheral analysis stage (Fig 1A; left panel) the model computes a time-frequency representation of the sound. This analysis resembles a typical short-time Fourier transform, but includes several features, more biologically plausible for the auditory periphery: this includes log-frequency cochlear-shaped filters, half-wave rectification and firing rate limitations.

During the Central analysis stage (Fig 1A; middle panel) the model computes a series of time-frequency analyses that vary in their spectral scale. These variations in scale capture the spectral dependency of measured receptive fields found in the inferior colliculus (IC) [69–72] and primary auditory cortex (A1) [64, 73, 74].

During the Object analysis stage (Fig 1A; right panel) the model computes a number of different scene interpretations. Each interpretation consists of one or more groupings of Central features into a stream or object. Consistent with the evidence concerning the formation of auditory objects in the brain, these features are grouped through a process of temporal coherence [8, 9, 65, 75, 76] and strung together over time on the basis of object continuity [9, 66, 77, 78].

We vary the behavior of these three stages across a large ensemble of variants by systematically varying the magnitude of three terms—adaptation, inhibition and noise (Fig 1B)—across the three analysis stages. These terms are applied to a set of weights which determine the relative strength of different frequencies (Peripheral), scales (Central) or scene interpretations (Object). Adaptation reduces each weight by a delayed, low-pass version of itself (self-referential link in Fig 1B). Inhibition reduces each weight by a delayed, low-pass version of neighboring weights (links to and from second unit in Fig 1B). Noise modulates each weight randomly (link to noisy response in Fig 1B). These basic components have been applied in various forms throughout the years to model perceptual bistability [18, 19, 36–38, 42–45].

### Model Evaluation

To evaluate each model in our ensemble we compare its ability to predict human behavior on two data sets, shown in dark blue in the panels of Fig 2; together these data sets capture two key hallmarks of perceptual bistability.

**Fig 2.**
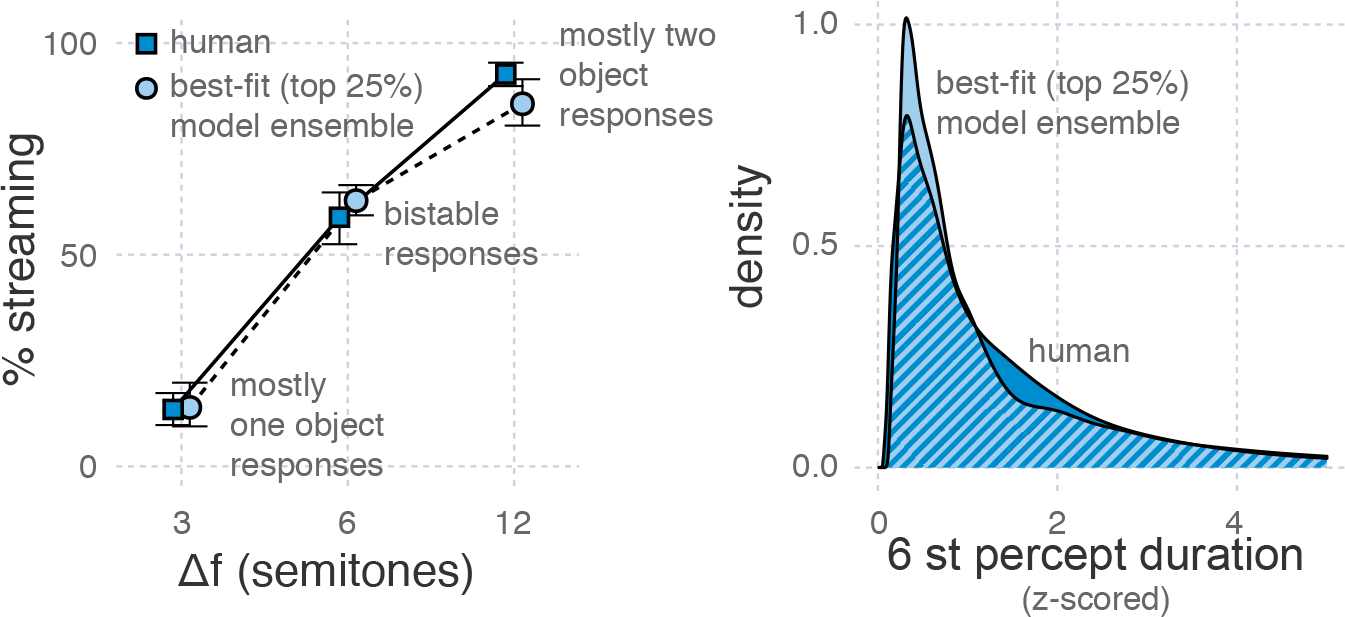
Best-fitting Models. The performance of humans and models that best fit human behavior for two key measures: the percent of streaming responses for each stimulus, and the distribution of response lengths. Models included here fell in the top 25^*th*^ percentile according to an aggregate measure of model fitness (the model:human deviation ratio). **A**. The percent of streaming (2 or more object) responses. Human data were taken from two existing studies [16, 79]. Error bars denote the 95% confidence intervals estimated by bootstrap. **B**. The distribution of response lengths for the 6 semitone stimulus. The lengths of each individual model and human listener were first z-scored in log space. Human data come from the control condition of an unpublished study.

The first hallmark of bistability is that it occurs only for specific, ambiguous stimulus configurations. Fig 2A shows human data which captures this hallmark [16, 79]. It depicts performance in terms of the percent of streaming responses (y-axis), across the three stimulus conditions tested (x-axis). A streaming response is one that indicates the listener heard two or more sounds, and a fused response indicates only one sound was heard. The pure tones in the ABA pattern presented to listeners were separated by either 3, 6 or 12 semitones (st). Only the 6 st stimulus elicited a bistable response: listeners reported the 3 st stimulus was mostly heard as a single sound source, and the 12 st stimulus as two streams of sound.

The second hallmark of bistability is the characteristic distribution of response lengths. The response length is the amount of time between when a listener reports hearing one stream and when they report hearing two streams of sound, and vice versa. Fig 2B shows human data which captures this hallmark (unpublished data, c.f. Human Data). It depicts performance in terms of the distribution (y-axis) of z-scored response lengths (x-axis), but only for an ambiguous stimulus. The figure shows that there were periods of relatively stable perception (the distribution has a long tail), with some characteristic length (the distribution peaks above zero). Z-scores for each individual configuration of the model and each human listener were computed in log space.

We chose to use different data sets across these panels because neither data set could capture the characteristics of the other: in the first two studies [16, 79] the trials were too short to avoid cutting off the tail of the distribution shown in Fig 2B, and so the pattern of response lengths could not be determined; the data presented in Fig 2B focused on bistable responses, using only the 6 st stimulus, and employed longer trials (unpublished data), and so there is no available data for the 3 or 12 st stimulus configurations shown in Figure 2A.

To evaluate each model’s ability to capture the first hallmark of bistability we are interested in—the selectivity of bistability—we compute a quantity referred to as the *response deviation*: the deviation of model predicted responses from that of the mean human listener. Specifically, for each stimulus (3, 6 and 12 st), we found the proportion of “streaming” responses (ala Fig 2) for each individual simulation run and for the average human listener, and then computed the root-mean-squared difference between model and human responses across all three stimuli.

To evaluate each model’s ability to capture the pattern of response lengths, we compute the *response-length deviation*: for each model simulation we compute the Kolmogorov-Smirnov statistic [80] of the model’s distribution of z-scored response lengths vs. that same distribution for the human listeners. We selected the Kolmogorov-Smirnov—the maximum difference between the empirical cumulative distribution function of two samples—for its sensitivity to subtle differences between two distributions.

These two measures of deviation are combined to provide an overall fitness score for each model, referred to as the *model:human deviation ratio*. To accomplish this, the two deviation measures are first placed on a comparable scale by dividing model deviation by the deviation of individual human listeners. The two deviation ratios are then averaged. We compute deviation for each individual human listener from the overall sample using the same procedure described above for computing model deviation. The larger the ratio, the greater the model deviation is relative to the deviation of the average human listener from the mean. A deviation of 1 would indicate the model has the same amount of deviation from the mean as the average individual human listener.

To determine whether our ensemble includes models that are consistent with human behavior, we examine the performance of the top 25% of models, according to this model:human deviation ratio (light blue circles in Fig 2A; light blue region in Fig 2B). These model data support the merits of our aggregate measure, confirming that when models score well (low) by this measure, they collectively show behavior quite similar to the mean human data.

### Model Behavior Across the Ensemble

We now evaluate the behavior of the entire model ensemble, using the aggregate model:human deviation ratio and the two separate deviation measures—response and response-length deviation (see Model Evaluation). We focus on the systematic variation of adaptation and inhibition both within and across the levels of the hierarchy. Varying the amount of noise across levels did not appear to change the key conclusions of our results (see Sensitivity Analysis).

Fig 3 shows the within-stage model variations of adaptation and inhibition: in these variations, adaptation, inhibition and noise are injected into only one of the levels of the hierarchy at a time, setting their parameters to zero for the other two stages.

**Fig 3.**
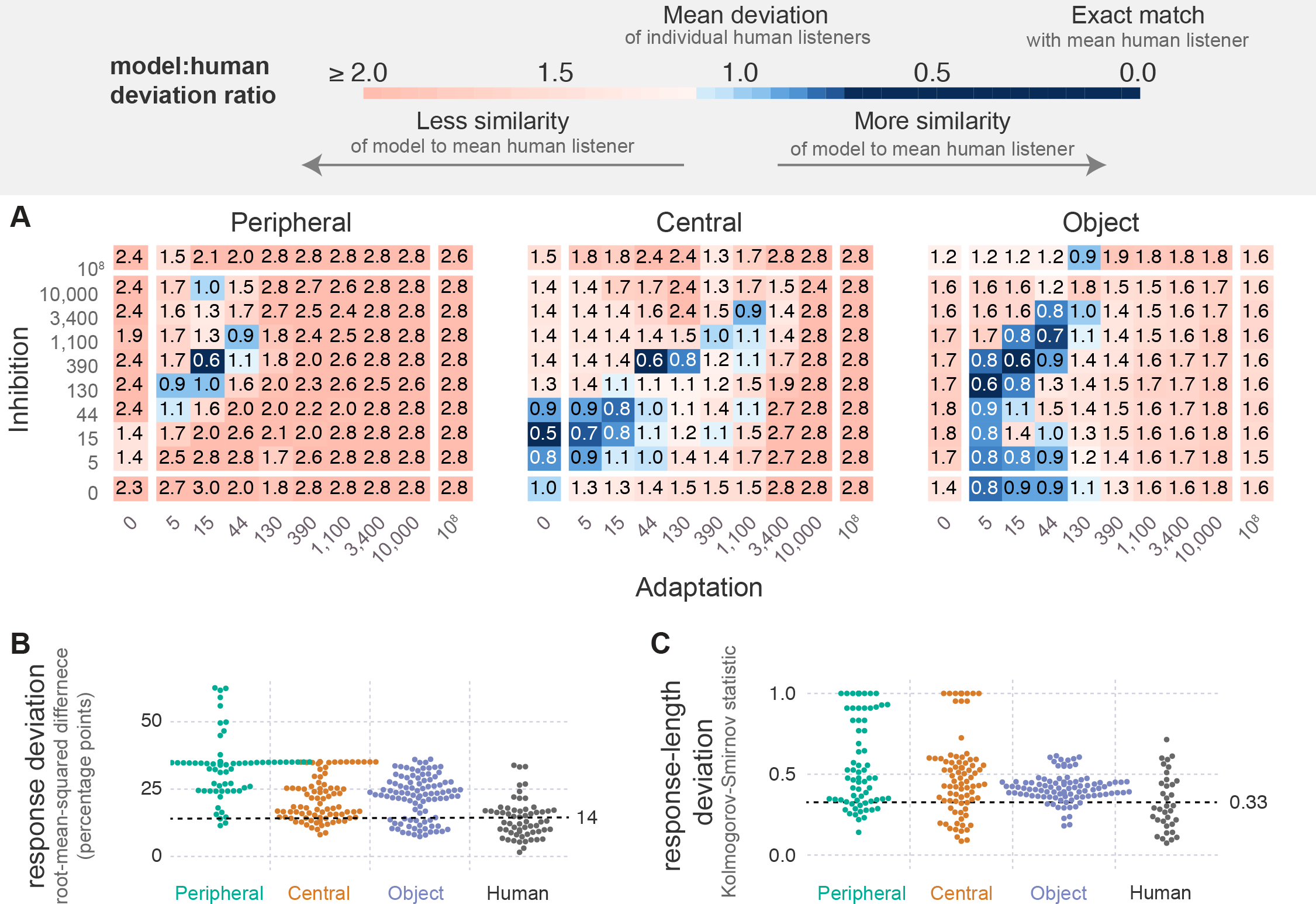
Within-stage variations of adaptation and inhibition. Each model shown includes terms for adaptation, inhibition and noise in one of the three analysis stages, leaving the other two stages with no such terms. **A** The model:human deviation ratio for all levels of inhibition and adaptation at each of the three analysis stages. Each square represents average deviation for all simulations of a single model from the ensemble with the given level of adaptation and inhibition. Values above one (pale blue to red) indicate models with more deviation than the average human listener; values below one (blue colors) indicate models with the same or less deviation than the average human listener. These measures are averaged across the ratio of both response deviation (panel C) and the response-length deviation (panel D). **B** The response deviation for all models and the human listeners from [16, 79]. Each point for the three model variations represents the average response deviation for all simulations of a single model from the ensemble. Offsets from the central y-axis of each condition are used as a visual aid, to ensure that all data points are at least partially visible. The dotted line marks the mean of the human data: the deviation ratio is greater than one above this line and less than one below this line. **C** The response-length deviation for all models and human listeners. Human data are from the same unpublished control condition shown in Figure 2. This panel follows the same format as described for panel B.

Fig 3A shows performance for the within-stage model variations in terms of the model:human deviation ratio. This measure is shown for each level of adaptation (x-axis) and inhibition (y-axis) tested within each of the three levels of the hierarchy (columns). The figure supports two key findings: that some variations in the parameters of each level lead to human-like model performance (there is blue in all three panels); and more such variations of the higher-level model parameters than lower-level parameters lead to human like performance (there is more blue moving from left to right).

The overall outcomes reflected by the model:human deviation ratio are also supported when we examine the two deviation measures separately: response deviation—in Fig 3B—and response-length deviation—in Fig 3C. Note that points are displaced along the y-axis within this figure to ensure that all datum are visible; thus, the overall shape of this plot provides an indication of the distribution of the data. Both figures show that some of the individual models (data points) fall at or below the mean human deviation (dotted lines). Further, as we move up along the hierarchy, the mass of points (the mean) is lower, meaning that there are increasingly more models more consistent with the human data.

Fig 4 summarizes the results of the across-stage variations of adaptation and inhibition: in these models, adaptation and inhibition are varied simultaneously across all three levels. Despite the presence of these terms across all three layers in many possible configurations, the results remain quite similar to the within-stage variations of Figure 3: there are non-zero levels of adaptation and inhibition at all three stages that lead to human-like performance (blue in all three panels), and there are more such accurate model variations as we move up the stages of analysis (blue increases from left to right). Note that each datum in these figures (square in A or point in B and C) represents the best (minimum) mean deviation ratio across all models with the given magnitude of adaptation and inhibition at a specific analysis stage. Therefore, for a given stage, each point is representative of the minimum deviation ratio of 625 models (5^2^ × 5^2^), one model for each level of adaptation and inhibition at the other two analysis stages.

**Fig 4.**
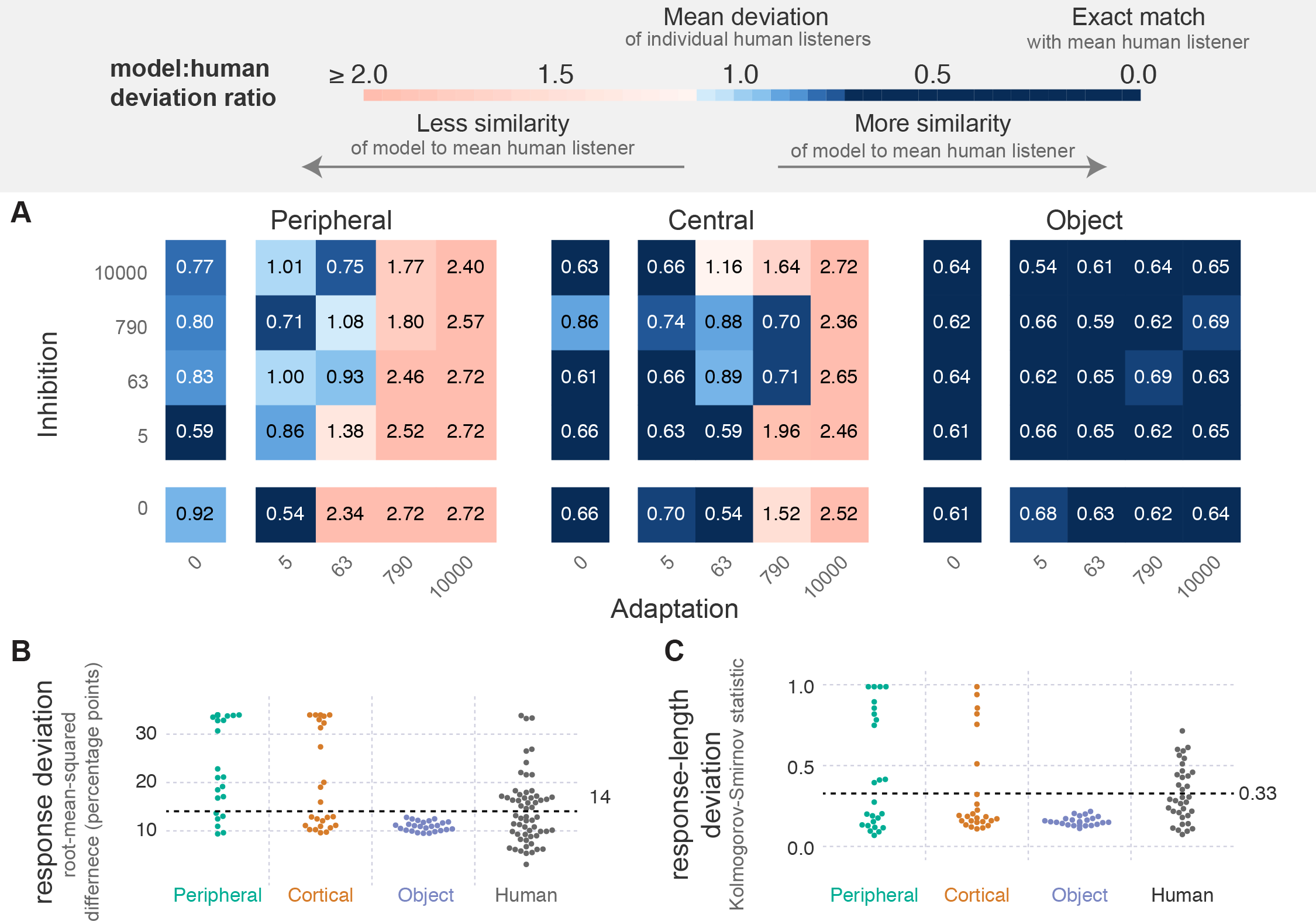
Across-stage variations of adaptation and inhibition. Each model in the ensemble includes some amount of adaptation, inhibition and noise at all three analysis stages. **A** The minimum model:human deviation ratio for each value of adaptation and inhibition at each level. Each square represents the average for all simulations of the best performing (minimum deviation) model with the given level of adaptation and inhibition. Values above one (pale blue and red colors) indicate models with more deviation than the average human listener, values below one (blue colors) indicate models with the same or less deviation than the average human listener. **B** Response deviation for the data shown in A: that is, the minimum percent-streaming deviation for each pairing of adaptation and inhibition, tested at each level of the model hierarchy. The dotted line indicates the average deviation of the human listeners. **C** Response-length deviation for the data shown in A. Figure follows the same format as described for panel B.

### Sensitivity Analysis

To determine how sensitive our results are to specific parameter values, we examine the effects of varying these parameters on the model:human deviation ratio (Fig 5). We vary the time constants for adaptation (*τ_a_*) and inhibition (*τ_b_*), the magnitude of noise (*c_σ_*), and the breadth of inhibition (Σ_*b*_). For each parameter considered we vary that parameter, while leaving all other parameters at their default value. For each parameter setting we examine the within-stage model variations, but using a coarser grid than shown for the within-stage model variations from Fig 3: instead we use the 25 parameter combinations shown for Fig 4. An inspection of the individual data revealed no large differences between the results, regardless of the parameter varied. (That is, we re-plotted Fig 3 for all 13 model variants)

**Fig 5.**
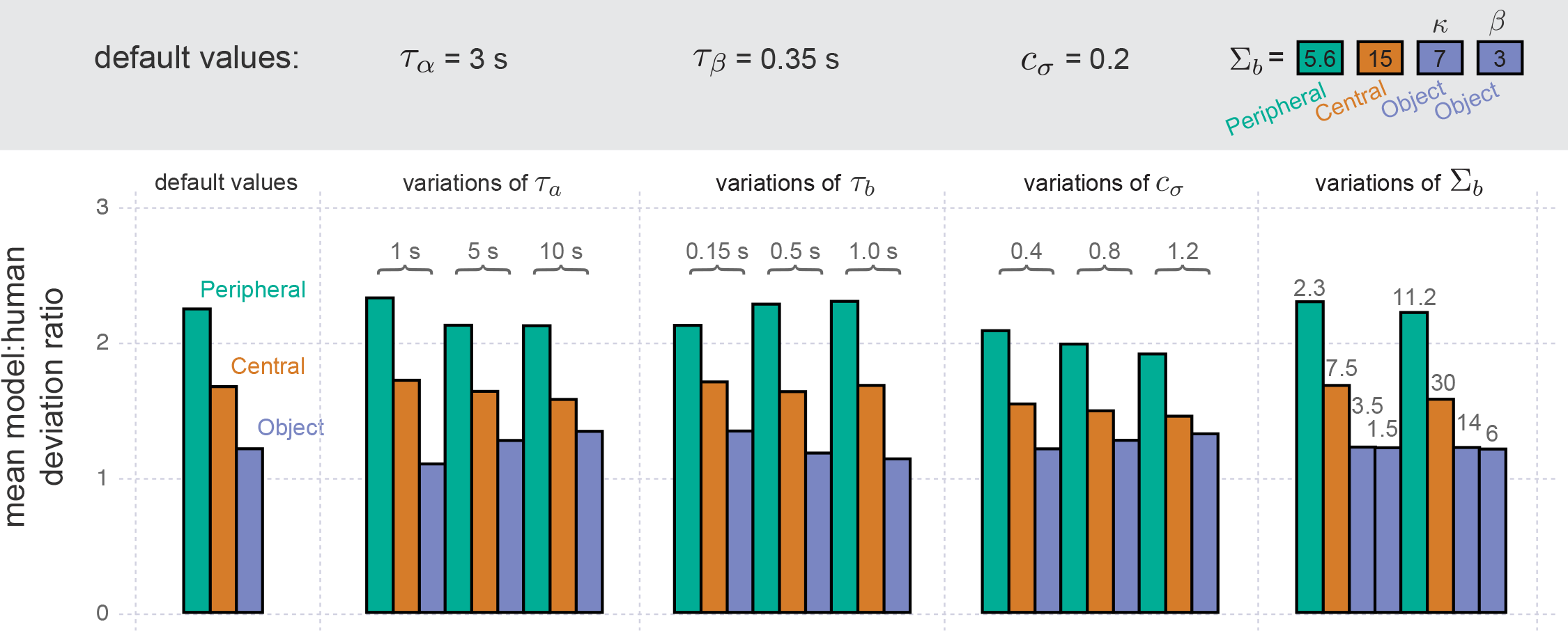
Sensitivity of model. A summary of the overall model performance across different parameter variations. The default-parameter model essentially summarizes the results from Fig 3A, though the models tested here use a coarser grid of variations across adaptation and inhibition, consistent with the grid used for the remaining parameter variations shown in the present figure. For each parameter varied (columns), we consider multiple possible alternatives (numbers above each group of bars), changing it from a default value (shown above each group, within the gray region). Note that the values of inhibition breadth (Σ_*b*_) differ across the levels because their effect on behavior differs across the levels.

Fig 5 provides a summary of all model variants. It shows the mean model:human deviation ratio (y-axis) for all the within-stage model variations when using the default model parameters (far left column) and when using a number of different parameter variants (x-axis). For ease of reference, the default model parameters are also shown (upper gray row). Note that the final parameter (Σ_*b*_) had distinct values within each level of the hierarchy because its meaning depends on the number of competing units, which varied across level.

This figure shows that one of the key results of our analysis is not altered by variations in the model parameters. Regardless of parameter variant, more variations of adaptation and inhibition are consistent with the human data as we move up the hierarchy. As such, the mean model:human deviation ratio across the variations of adaptation and inhibition decreases as we move up the hierarchy.

Furthermore, all of these parameter variants have some model variations consistent with the human data. The average minimum model:human deviation ratio across these variants is 0.88 (SD = 0.18).

## Discussion

In the model ensemble reported here, each model includes three levels of processing—Peripheral (Fig 1B; left panel), Central (Fig 1B; middle panel) and Object (Fig 1B; right panel). We systematically vary the amount of adaptation, *a*(*t*), inhibition, *b*(*t*) (Fig 1B) and noise, *n*(*t*), within each level separately (Fig 3), and across all levels simultaneously (Fig 4). We then compare model performance to past reports of human responses to a pure-tone ABA pattern; human listeners were asked to respond continuously, indicating whether they heard a “fused” (1 object) or “streaming” percept (2 or more objects).

There are two key findings: (1) there are models consistent with human behavior regardless of the locus of bistability we consider here, either within each analysis stage (Fig 3 shows blue in all three panels) or across all three simultaneously (Fig 4 shows blue in all three panels); (2) the number of variations of adaptation and inhibition generating plausible bistability increases as the locus of bistability within the perceptual hierarchy increases. This increase in viable model variation up the hierarchy is true both for the within-stage variations (Fig 3; more blue to the right) and for the across-stage variations (Fig 4; more blue ot the right). Note that the standard for “plausible” in this case is that a model only generate perceptual bistability for one (6 st) of the three stimuli presented to the model (3, 6 or 12 st; Fig 2A) and that the pattern of perceptual switching follows the systematic, log-normal-like distribution characteristic of perceptual bistability (Fig 2B). These results appear relatively robust, as they are not substantially altered by variations of several model parameters (Fig 5).

These findings are especially interesting in light of the existing evidence that there are substantial individual differences in perceptual bistability [21–29]. The implication is that (1) perceptual bistability is pervasive and can emerge across many levels of the auditory hierarchy and (2) there may be greater individual variation in the higher level stages of auditory scene analysis, and less variation at the earliest stages, at least in terms of the magnitude of adaptation and inhibition.

The notion that perceptual bistability may emerge from many different stages of analysis has received converging support from a number of different sources [3, 10, 12–14, 16–20, 43, 45–58]. However, existing efforts to model the locus of bistability across a hierarchy [11, 13, 14, 17, 36] differ from the present effort in that they (1) do not consider multiple possible loci of bistability within this hierarchy and (2) do not introduce adaptation and inhibition within a system that actually computes a scene interpretation from raw sensory input. These two differences strengthen the support for the pervasiveness of perceptual bistability because they show that (1) bistability can emerge from a large variety of different model configurations and that (2) this emergence can scale from simple mathematical models to a more complete system capable of generating perceptual inferences relevant to behavior. Outside the region of an ambiguous stimulus presented to our model (6 st), the model computes a useful mask that either fuses a scene into one sound (for 3 st) or extracts the stimuli into two separate sound streams (12 st).

Past computational studies of bistability during auditory streaming have focused on the role of a single locus of perceptual bistability at either relatively early stages [60, 61] or later stages of processing [59, 81]. None of them systematically vary the locus of perceptual bistability. In Rinzel et al [60] (and also [61]), a relatively low-level locus of auditory bistability is examined: three broadly tuned frequency channels compete and these can be used to explain bistability during the ABA tone pattern. This approach is similar in spirit to introducing bistability during the Central or Peripheral analysis stage of our model. In Mill et al [59] and in Barniv and Nelson [81] a higher-level locus is evaluated, in which probabilistic interpretations of the acoustic scene compete with one another. The representation used assumes a previously processed, discrete set of events. The implicit assumption is that bistability occurs over some representation of auditory objects. In this way it is similar to introducing bistability during the Object stage of our model. These existing models of auditory streaming represent input in a relatively abstract form, that differs across the models, limiting the ability to compare the merits of the two different loci within a single system. Existing auditory models of this auditory streaming paradigm that do include a more detailed perceptual hierarchy do not appear to demonstrate any form of perceptual bistability [**?**, 65, 76]

The existing evidence for individual differences in perceptual bistability suggest that the breadth of functional variations represented by our model ensemble may also be present in the human population. There are three types of outcomes that support the notion that the mechanisms of bistability vary across listeners. First, there are behavioral reports of individual variability of bistability [22, 82, 83]. These behavioral differences lead to patterns of perceptual switching that differ more across listeners than across sessions of the same listener [22]. Second, these switching patterns can be accounted for using differences in the dynamics of adaptation, inhibition and noise [5, 7, 18, 29, 38, 42–45, 84]. Third, there are a number of neural correlates of these behavioral signatures in the balance of excitation and inhibition, as measured by magnetic resonance spectroscopy (MRS) [27, 29, 85].

If we accept this premise, it implies that adaptation and inhibition in the human auditory system may be more precisely tuned at earlier stages of analysis and have much greater variation at higher stages of analysis across individuals. The prediction that there is greater variability in higher stages of analysis is consistent with three existing lines of evidence. First, it is consistent with the overall trend from a number of studies of resting state activity [86, 87], showing a coarser outcome: they reveal greater variability in regions of the brain associated with more cognitive processing, than regions associated with perceptual processing. Second, it is consistent with studies of perceptual bistability indicating that there may be greater individual variability in the balance of excitation and inhibition in regions associated with more cognitive processing [27, 29, 85]. Third, it is consistent with the prevalence of evidence supporting the role of more general, higher-level processing stages during perceptual bistability [4, 7, 20, 33, 51–58]; if these higher-level sources play a more dominant role in perceptual bistability, this could be due to the greater variety of configurations at later stages of analysis that can generate perceptual bistability. While consistent with these prior reports, to our knowledge this study is the first to specifically suggest that individual variability may increase from peripheral to more centrally computed features, and from these features to object-level processing. It is precisely our use of an explicit ensemble of models with varying levels of adaptation and inhibition across these specific stages of processing—a configuration apparently unique to our particular modeling approach—which generated this evidence.

What specific properties of the model lead to greater variations at higher stages of analysis? The most fundamental change across levels is how a stimulus is represented, and seems the most plausible account. This representation fundamentally changes what exactly the oscillatory activity generated by adaptation and inhibition operates over: frequencies (Peripheral), frequency scales (Central) or scene interpretations (Object). The case for some fundamental property of the stimulus representation being important is further strengthened by our sensitivity analysis, which showed no change in the relative advantage of the higher stages of analysis over lower stages across a number of different parameter configurations (Fig 5).

If there are indeed more variations of object-level bistability than earlier stages of processing, one explanation of this difference in variation would be that the earlier stages of analysis have been optimized by experience and evolution to remain relatively stable, while higher-level stages are more adaptive and flexible. This would be consistent with the evidence that later stages of processing are subject to more explicit, consciously driven forms of plasticity [34, 88–90] and encode more idiosyncratic properties of stimuli, that vary with subject expertise [91–94].

The present model also takes a step in the effort towards scaling the intuitions developed with simpler, more controlled models to a larger system capable of actually computing behaviorally relevant information from raw sensory data. This exercise in scaling is important for at least two reasons. First, because the model can take as input any auditory signal, it opens the door to future efforts to integrate our understanding of auditory bistability across multiple paradigms. Human listeners show apparent bistable responses to a number of other auditory stimuli, including Shepard tones [95–97] and repeated words [6]; natural extensions of our Central and Object stages of processing could be used to model these additional tasks. Second, it provides a useful test of simpler, more abstracted models of adaptation, inhibition and noise [18, 19, 38, 42, 43, 45]. If our theories of brain function are accurate, then it should be possible to replicate the behavior generated by the brain with these theories. The present model contributes to this effort by showing how adaptation, inhibition and noise may generate auditory bistability within a more comprehensive system. Past efforts that may contribute to our understanding of how brain-like computations can scale to larger systems fall short of this demonstration either because they include comprehensive dynamics of adaptation, inhibition and noise but abstract away from the specific challenges of auditory scene analysis [98–101] or because they can perform some form of auditory scene analysis but do not show any identified form of adaptation, inhibition and noise nor any bistable output [102–104]. Future efforts towards scaling of the present model could build on the strengths of any one of these past systems.

The present study compares three different forms of bistable competition across different stages of auditory processing, but there are at least two additional possible forms of bistable competition that might be considered: there are indications that (1) there is competition across different levels of the perceptual hierarchy [5, 7], and (2) there is a hierarchy of bistable oscillations, with more cognitive bistable oscillations modulating the rate of a more rapid oscillation at a perceptual level [105]. Given the ability of the three forms of competition examined in the present model to co-exist within a single system, we speculate that additional forms of competition may co-exist to varying degrees. At the very least the work here suggests that that it would be fruitful to systematically compare these proposed sources of perceptual bistability within a single system.

Bistable perception has proven to be a potential means for examining the differences between conscious and unconscious states [33–36], contrasting cases where listeners are conscious or unconscious of particular perceptual interpretations. The present model may thus provide a potential baseline upon which to develop computational models of conscious vs. unconscious perception in auditory scenes. Two tentative conclusions from the study of consciousness are relevant to interpreting the present model: first, it appears that the neural correlates of consciousness generally display a relatively long latency on the order of several hundred milliseconds [33, 34, 106] (but see [76]). Second, these conscious states appear to be associated with more comprehensive integration of information [34, 106]. This evidence favors an interpretation in which the present model largely represents unconscious processing, given the early stages of the auditory system it may correspond with and its feed-forward nature. The additional sources of bistability such as top-down, predictive error signals [5, 7] could have more of a role to play in any consciously driven aspects of perceptual bistability [34, 76].

## Conclusion

The present computational study indicates that auditory bistability can plausibly arise from a multitude of different sources, from very early following the auditory periphery to later, object-level encodings of the stimulus. Given the pervasiveness of adaptation, inhibition and, arguably [100], noise, throughout the nervous system, this suggests that bistability arises from these sources throughout the auditory hierarchy. The results also indicate that there are more variations of the later stages consistent with human data than earlier stages of processing. When interpreted in light of the significant individual differences found in perceptual bistability [21–29, 85], a possible implication is that there is greater variability in human function for the higher-level stages of auditory processing, than lower-level stages of processing.

## Materials and methods

### Model Design

In each model in the ensemble the three stages of analysis are a Peripheral analysis, *P* (*t*), a Central analysis, *C*(*t*), and an Object analysis, *O*(*t*)—shown in Fig 1A. Into each level we inject adaptation, *a*(*t*), inhibition, *b*(*t*), and noise, *n*(*t*)—collectively denoted *F* [*x*(*t*)], shown in Fig 1A—with varying magnitudes. All model parameters are described in Table 1 and 2.

**Table 1.** All model parameters, some of which took on multiple values; multiple, consecutive integers are denoted with “ … ”.

|  | value | description | Eqs |
| --- | --- | --- | --- |
| $f$ | $440\text{Hz} \times 2^{\frac{-1\dots17-0.5}{12}}$ | frequency channels | 1–4 |
| $\omega$ | $2^{-1\dots2}$ cycles/octave | frequency-scale channels | 5–7 |
| $\psi$ | $2^{1\dots5}\text{Hz}$ | temporal-rate channels | 8 |
| $r$ | 2 | components of NMF factorization | 9 |
| $T$ | 30 | Object-level history | 11 |
| $\sigma_F$ | 5 log Hz | object continuity neighborhood | 12 |
| $\kappa$ | $15s/\Delta t, 20s/\Delta t$ | source prior, mean degrees of freedom | 13 |
| $\alpha$ | 1 | source prior, shape for variance | 14 |
| $\beta$ | 0.25, 0.5, 0.8 | source prior, scale for variance | 14 |
| $Z_\alpha, Z_\beta$ | 2, 2 | bernouli prior for object presence | 16 |
| $c_a$ | values shown in Fig 3–4 | magnitude of adaptation | 19 |
| $\tau_a$ | 3 s | time constant of adaptation | 20 |
| $c_b$ | values shown in Fig 3–4 | magnitude of inhibition | 21 |
| $\tau_b$ | 0.35 s | time constant of inhibition | 22 |
| $\theta_b$ | 6 | strength of inhibition neighborhood | 23 |
| $c_n$ | 0.2 | magnitude of noise | 24 |
| $\tau_n$ | 500 ms | time constant of noise | 24 |
| $\Theta_r$ | 0.75 | heuristic threshold ratio | 29 |
| $\Theta_c$ | 0.996 | heuristic global threshold | 29 |
| $\Theta_f$ | 0.95 | heuristic local threshold | 29 |
| $\Theta_w$ | 500 ms | heuristic time window | 29 |
| $\Theta_\Delta$ | 250 ms | heuristic time step | 29 |

**Table 2.**
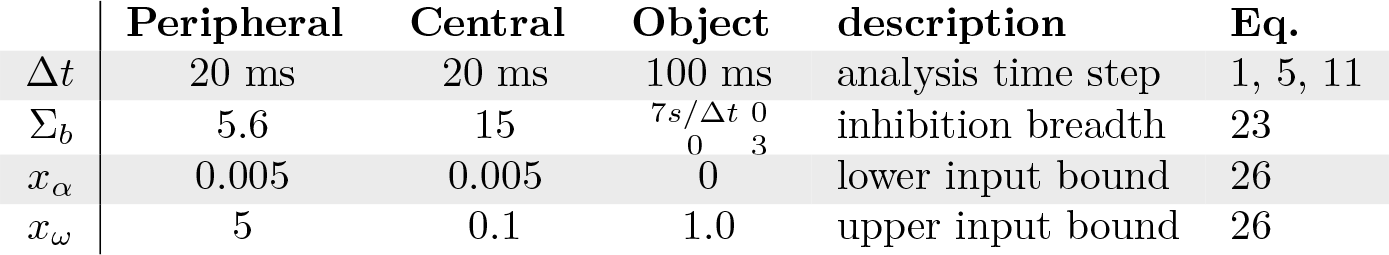
Model parameters that varied by level of analysis.

In the below formulas, we use superscripts to denote a variant of a function (e.g. *P* vs. *P^F^*), and subscripts to denote slices along a dimension implied by the variable name (e.g. *P*_*f*_ (*t*) is a slice along the dimension of frequency). A slice along *f* includes all values of the function at time *t* such that frequency is equal to *f*. Note that the absence of a subscript implies all values are included (e.g. *P*_*f*_ (*t*) is a slice of the function along frequency and *P* (*t*) includes all frequencies).

### Analysis Stages

The three analysis stages of each model in the ensemble are a Peripheral (Fig. 1A; left panel), Central (middle panel) and Object (right panel) analysis. Each analysis stage is computed for a series of time slices separated by ∆*t* (c.f. Table 2)

The Peripheral analysis (Fig. 1A; left panel) reflects processing that occurs shortly after (e.g. Cochlear Nucleus) and at the periphery of the auditory system as described in [64]. It is similar in concept to a log-frequency spectrogram, but aims for a more physiologically realistic functional form: first, cochlea-like filter shapes *h*_*f*_ are applied to the original time-amplitude signal *x*(*t*) (Eq 1), followed by local inhibition (Eqs 2; with 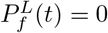 for *f* = 0). This is followed by half-wave rectification []^+^, and a low pass filtering of each frequency channel, *h*_*lp*_ (Eq 3). Finally, adaptation, inhibition and noise (*F* [*x*]) are applied along the frequency channels *f* (Eqs 4).

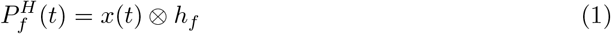

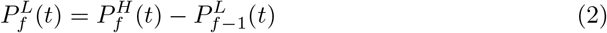

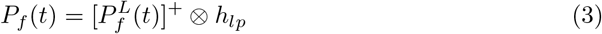

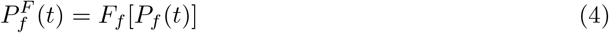

The Central analysis (Fig. 1, middle panel) reflects processing that occurs in the midbrain [69–72] and primary auditory cortex (A1) [64, 73, 74]. There are quite a number of proposed feature dimensions (e.g. pitch [107], location [108]) processed sometime after the initial peripheral analysis. Here, given the nature of our stimulus and task, we focus on a single dimension: spectral scale. Following Chi et al [64], spectral scale is captured via a wavelet transform of the auditory spectrogram along the frequency axis. The result is a sequence of complex-valued spectrograms (Eq 5), each reflecting the amplitude and phase response to the spectrogram at a different spectral scale *ω* (in Fig 1, middle panel, the amplitude of these responses are shown). Adaptation, inhibition and noise (*F* [*x*]) are then applied by first computing the average magnitude for each scale (Eq 6), and rescaling by these magnitudes (Eq 7).

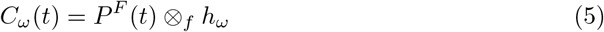

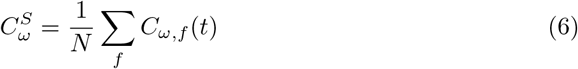

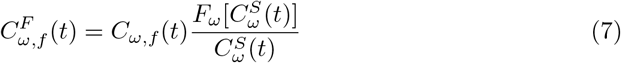

In Eq 6, *N* is the number of frequency bins (c.f. Table 1, first row).

The Object analysis reflects computations that occur during the segmentation of sounds into objects (also called streams): it computes multiple, probabilistic groupings of the features in a sound scene and selects one of them (Fig. 1B; right panel). It is a novel implementation, similar in spirit to some existing, more abstract models of probabilistic scene interpretation [59, 81]. Its output is a series of time-frequency spectrograms of each sound source. Two basic principles guide the grouping of features into objects: first, we employ temporal coherence [**?**, 65, 75, 76]—the notion that features which change at the same rate likely originate from the same object—second, we employ object-continuity [9, 66, 77, 78]—the notion that, all else being equal, features originating from the same object tend to change smoothly in time. There are two stages of computation: a near-simultaneous and a sequential grouping stage.

In the first stage of computation, a set of near-simultaneous groupings of features are found by using the principle of temporal coherence. Specifically, a set of components, *K*(*t*), are found using a non-negative factorization of each time window *w*(*t*) across many temporal rates of the features of *C*(*t*), the output of the Central analysis. This factorization of different temporal rates allow us to compute features that consistently change in a synchronous manner across the window of analysis *w*(*t*). Specifically, the components *K*(*t*) are defined by Eq 8–9.

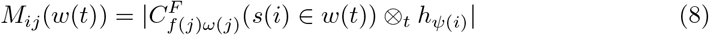

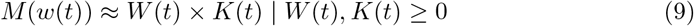

The matrices *W* and *K* at each time *t* are low-rank approximations of *M* with inner dimension *r*. Each column (*j*) of matrix *M* represents the features of *C^F^* (*t*): frequencies are selected by *f* (*j*) and scales by *ω*(*j*). Each row (*i*) of *M* represents a temporal rate of change, *ψ*(*i*), at a given time slice, *s*(*i*), within a small window of time, *w*(*t*). The rates of change are selected by rate filters *h_ψ_*, which are convolved along the time axis (⊗_*t*_) with *C*(*t*). These filters are analogous to the spectral filters used to compute *C*(*t*): they each extract a distinct rate of change (amplitude modulation), and are described in [64]. We solve Eq 9 using the algorithm from [109].

In the second stage of computation, a sequential grouping of the components is found by applying the principle of object continuity. Specifically, for each interpretation *i*, components *K*(*t*) are organized into a set of sources *g*_*i*_(*t*), all some of which may be present at a given time (*t*). Each sources *h* is defined by the sum of its constituent components, at every time slice *t* (Eq 10). In our simulations we consider up to *r* sources (c.f. Table 1).

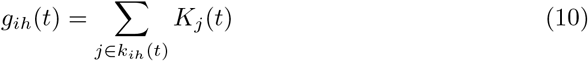

We solve for the set of components *k*_*ih*_(*t*) belonging to each source *h* according to Eq 10–15 using a simple greedy approach: we compute the solution for each time step *t* sequentially, holding all prior time steps constant. The grouping of components for all sources *g*_*i*_(*t*) constitutes a single interpretation, *i*. We found one such interpretation for each pair of values for the hyper-parameters *κ* and *β* (c.f. Table 1); these hyper-parameters determined the degree of feature smoothness assumed by a given interpretation and their role is described below, more precisely.

We model the observation of a source at time *t* as a multivariate Normal distribution consistent with the last *T* observations of that source (Eq 11).

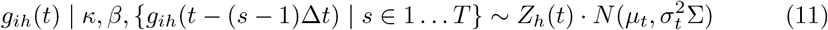

∆*t* is the time step (c.f. Table 1). The integer *Z* (0 or 1) reflects the fact that not all sources would be present at all moments in time. The correlation matrix Σ is key to encoding the principle of object continuity. It is a fixed matrix that had greater correlation when the frequency *f* of component *i* was closer to the frequency of component *j* (Eq 12).

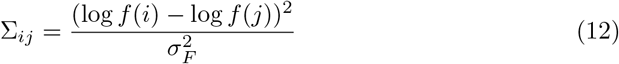

This correlation structure therefore encodes the principle that objects will smoothly transition along their frequency components (*σ_F_* is defined in Table 1).

The distribution of *µ* and *σ* determine the strength of the assumption of object continuity, and are conjugate priors for ease of computation (a Normal distribution in Eq 13, and Gamma distribution in Eq 14). The shape of their distributions depends on two variable hyper-parameters: the scale of prior mean, *κ*, and the scale of the variance, *β* (*α* is found in Table 1).

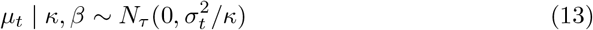

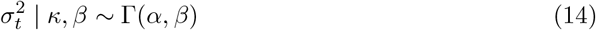

As noted above, the variable *Z* (Eq 15) determines the presence (1) or absence (0) of each object within a given frame. It has a Bernouli distribution, and is given a conjugate prior (a Beta distribution) for ease of computation, with fixed parameters *Z_α_* and *Z_β_* (Eq 16)

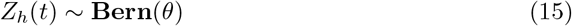

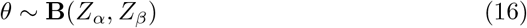

Finally, the dominant interpretation at each time frame (Eq 17) is used to compute the time-frequency mask of *O*(*t*) (Eq 18). Adaptation, inhibition and noise (*F* [*x*]) are applied to the relative log-probabilities of each interpretation.

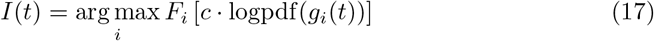

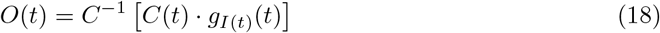

The constant *c* is selected to normalize the log probability distribution function (logpdf) such that the largest value after the first second of output was 1 (the initial logpdf values can be quite erratic, so we avoid values in this first segment). The function *C^−^*^1^ denotes the inverse of *C*(*t*) and its output is a time-frequency representation in the same format as the Peripheral analysis *P* (*t*).

### Adaptation, Inhibition and Noise

Adaptation, inhibition and noise follow a similar functional form as found in [43, 60], and are applied to a set of input weights *x*(*t*) within each stage of analysis, yielding a new set of weights *y*(*t*) (Fig. 1B).

Each term modulates the unsmoothed weights *y^U^* (*t*) and is characterized by a magnitude *c*—which determines how much the term modulates the output—and a time constant *τ* —which determines how quickly the term changes.

The amount of adaptation for each weight 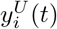 is determined by a low-pass, delayed version of 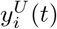 (denoted *a^D^*), shown in Eqs 19–20

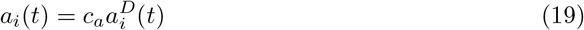

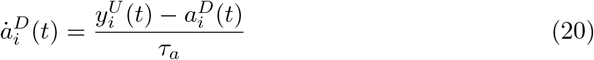

The dot (̇) above a function is used to denote the first derivative of that function. All functions are solved using a first order, finite-differences approach, using a delta specific to the particular stage of analysis (see Table 2).

The amount of inhibition for each weight 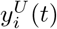 is determined by a low-pass, delayed version of distant neighboring weights, shown in Eq 21–22. The neighbors are selected by column vector *B*_*i*_ of weight matrix *B*, which varied depending on the level of an analysis.

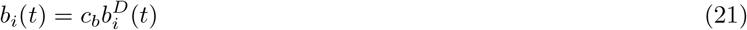

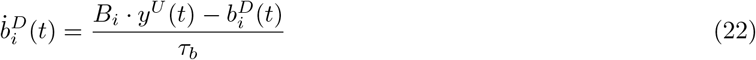

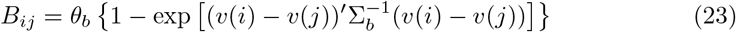

Each value in the inhibition weight matrix, *B*_*ij*_, is proportional to the distance between the two values: for the Peripheral analysis this is a scalar, in log frequency, for the Central it is a scalar in log cycles per octave, and for the Object analysis it is a two dimensional vector, the first term indicating the number of samples of prior data (for *κ*, c.f. Eq 13) and a scale value (for *β*, c.f. Eq 14). These distances are scaled by the distance weighting matrix Σ_*b*_ (c.f. Table 2).

The amount of noise for each weight 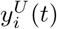 is defined by *n*_*i*_(*t*) (Eq 24)

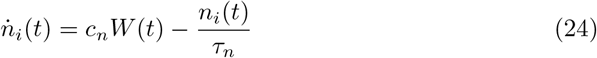

The *W* (*t*) term is a Wiener process (a.k.a. Brownian motion).

The three terms are used to modulate a bounded, smoothed version of the input weights (Eq 26) resulting in the initial, unsmoothed output of each weight 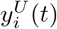 (Eq 27).

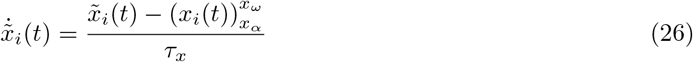

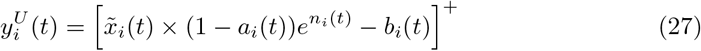

In the above equations, [ ]^+^ denotes half-wave rectification (all values below zero are set to 0), and 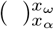 indicates a soft sigmoid bound between *x_ω_* and *x_α_*.

The final output weights are then computed according to Eq 28.

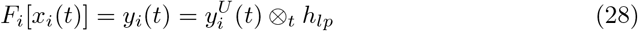

The ⊗_*t*_ denotes convolution in time with *h*_*lp*_, a low-pass Butterworth filter of order 3 at 1.5Hz. The low-pass filter ensured that brief changes in amplitude (due to the silence between tones) did not dramatically change the overall effect of adaptation and inhibition.

### Model Evaluation

We evaluate model responses on an ABA stimulus, created to exactly match the stimulus used in Snyder et al [16]. It consists of two different frequencies: one at 500 Hz (A) and another either 3, 6 or 12 semitones (st) above 500 Hz (B). Each triplet consisted of three equally spaced 50 ms tones, with onsets separated by 120 ms. The triplets are separated by 480 ms each. Pure tones included a 10 ms cosine ramp at the onset and offset.

For each set of the within-stage model parameters evaluated, we run 20 simulations, each with a 48 second stimulus composed of 100 repetitions of the ABA pattern. This is repeated three times, for the three stimulus conditions (3, 6 or 12 st). For each of the across-stage model variations, we used the same procedure, but it terminated early for all poorly performing model variations (Fig 4): following the first 10 simulations of the 6 semitone stimulus (always the first stimulus evaluated), if the model:human deviation ratio is greater than one, no more simulations for that model are run. This procedure ensures that our finite computational resources were not spent evaluating the merits of models that clearly performed poorly on this most essential stimulus condition.

To interpret the model output for each simulation, we use the following strategy. During the streaming task human listeners are often asked to respond continuously, reporting if they heard a “fused” (1 stream) or “streaming” percept (2 or more streams). To obtain a similar response for the model we interpret the maximum amplitude time-frequency mask found at each frame of *O*(*t*) as a “fused” or “streaming” response using an automated heuristic. Specifically, we compare the estimated frequency bandwidth of this dominant mask to that of the input’s time-frequency representation, *P* (*t*), and reported a 2 stream percept only if this ratio was greater than Θ_*r*_ (c.f. Table 1).

To estimate the frequency bandwidth, we use the following procedure. The frequency bandwidth of the dominant source is found for multiple overlapping windows of analysis *w*_*b*_(*t*). For each time window (of length Θ_*w*_, at time step Θ_∆_) this bandwidth is estimated by finding the maximum distance between frequency channels whose 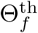 quantile falls above a data-driven threshold, determined by the 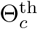 quantile of the entire mask. Specifically, using *Q*_*f*_ (*p*) to denote the *p^th^* quantile of frequency channel *f*, we estimate the bandwidth as follows

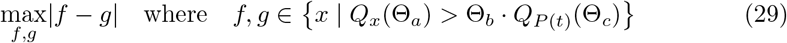

The values for all Θ parameters (shown in Table 1) are selected by hand to provide model responses consistent with a visual inspection of sampled output masks. We eliminate any brief switches: any response that returns back to the original response within 250 ms is discarded as a spurious output of our decision making heuristic.

Before analyzing the model responses computed with this procedure, we first discard the first response, *unless* it is longer than 9.6 seconds. Removing the first stimulus is consistent with the way human data are generally analyzed to avoid the “buildup” period [4]. However, we want to avoid ignoring responses during a substantial portion of the simulation time, and selected 9.6 as 20% of the full 48 second stimulus length. There are a number of poorly fitting models for which this first stimulus constitutes a substantial majority (> 90%) of the total stimulus length; dropping the first stimulus in these cases would mean the analyzed data would not be representative of those simulations.

### Human Data

We use human data from three sources: [16] and [79]and the control condition of yet-to-be published data. From the two published data sources, we use the context phase of Experiment 1A of Snyder et al [16] and the context phase of Experiment 1 and 2 of Yerkes et al [79]. The stimuli are identical to those described above, but the base frequency was 300 Hz instead of 500 Hz in some of the conditions (the measures of interest here did not appear to differ across these two different frequencies). We found the proportion of streaming responses following the stimulus build-up period by selecting response between the time range of 4 to 6 seconds.

The previously unpublished control data included a total of 35 normal-hearing participants (22 females) with an average age of 24.5. Listeners were recruited from the University of Nevada, Las Vegas community and paid for their participation. All procedures were approved by the University of Nevada, Las Vegas Office of Research Integrity. During the condition analyzed here, participants were presented the stimulus described above with a base frequency of 400 Hz, presented over 3M E-A-RTONE insert ear phones in a sound attenuated booth, at a level of 60 dB. Each trial of the stimulus was 48 seconds long, consisting of a total of 100 repetitions of the ABA sequence. For the duration of each trial, listeners were asked to continuously report whether they heard 1- or 2-stream percepts, with a short practice session during which the galloping rhythm of the 1-stream percept, and the more steady rhythm of the 2-stream percept were explained.

### Statistical Analyses

All reported confidence intervals are computed using a non-parametric bootstrap [110] with 10,000 samples. The density curves shown in Fig 2B were computed using a method for log-space density estimation [111] with a Normal kernel, where kernel bandwidths were determined by the Silverman rule [112].

### Code availability

The software for this study—made possible by the Julia programming language [113]—can be downloaded from https://engineering.jhu.edu/lcap/index.php?id=downloads or https://github.com/haberdashPI/bistable/tree/paper_submission

## Acknowledgments

This research was supported by Office of Naval Research grants N000141612879, N000141912014 and N000141712736 and National Institutes of Health grants T32DC000023, U01AG058532 and R01HL133043. The funders had no role in study design, data collection and analysis, decision to publish, or preparation of the manuscript. We thank Nathan C. Higgins and Brenne D. Yerkes for assistance with the behavioral data sets.

